# Mechanics and buckling of biopolymeric shells and cell nuclei

**DOI:** 10.1101/197566

**Authors:** Edward J. Banigan, Andrew D. Stephens, John F. Marko

## Abstract

We study a Brownian dynamics simulation model of a biopolymeric shell deformed by axial forces exerted at opposing poles. The model exhibits two distinct linear force-extension regimes, with the response to small tensions governed by linear elasticity and the response to large tensions governed by an effective spring constant that scales with radius as *R*^−0.25^. When extended beyond the initial linear elastic regime, the shell undergoes a hysteretic, temperature-dependent buckling transition. We experimentally observe this buckling transition by stretching and imaging the lamina of isolated cell nuclei. Furthermore, the interior contents of the shell can alter mechanical response and buckling, which we show by simulating a model for the nucleus that quantitatively agrees with our micromanipulation experiments stretching individual nuclei.

## INTRODUCTION

Polymeric shells are ubiquitous in biological and biomimetic systems, and they provide structure and mechanical robustness in objects as varied as pollen grains (1), viruses (2, 3), bacteria (4), protein and lipid vesicles (5–13), red blood cells (14, 15), and cell nuclei (16–19). These structures are composed of a thin layer of repeated monomeric and/or filamentous subunits organized in a spherical, ellipsoidal, or polyhedral geometry. They are often found in highly active biological (or biomimetic) environments in which they must resist and respond to perturbations, such as *∼* nN forces inside living cells.

The cell nucleus provides a particularly important example of a biopolymeric shell, the nuclear lamina. The lamina is a randomly connected polymer network of lamin intermediate filament proteins that resides at the periphery of the nucleus beneath the nuclear envelope and encloses the genome and other nuclear contents (18, 20). Experiments have shown that the lamina, particularly the lamin A/C protein, is an important component of cell nuclear elasticity, resisting external forces of 1 ≳ nN (16, 17, 19, 21–26). Defects in the lamina are associated with shape abnormalities known as blebs, which occur in conditions such as progeria, cancer, heart disease, and muscular dystrophy (27). These deformations of the lamina are hypothesized to arise via defective force response (28–32). Indeed, two models have investigated this hypothesis, finding that phase separation between different types of lamins (33) or disruption of the connectivity of the lamina (34) can result in bleb-like deformations of shells. However, recent experiments demonstrate that chromatin, the packaged genome inside the nucleus, also controls cell nuclear mechanics and morphology, and in some cases, is the dominant component (19, 35–37). To understand this interplay, it is critical to understand the mechanics and shapes of polymeric shells subjected to external forces.

The mechanics and morphology of polymeric (and elastic) shells have previously been studied with continuum elastic and statistical mechanical theory and simulations, as well as in several experimental systems. When indented by small point forces, shells exhibit a linear elastic response (12, 38–42). Larger forces induce a nonlinear response (38–42), and under sufficient loads, shells buckle inward, relieving the cost of stretching by forming localized bends (12, 39, 43, 44). Buckling can also be induced by pressurization of the exterior environment; above a critical external pressure the shell collapses, forming large scale folds and/or facets to reduce the enclosed volume (42, 45–47). However, less is known about the mechanical and morphological response of polymeric shells to large extensional deformations.

To understand the mechanics and morphology of biopolymeric structures such as cell nuclei, we developed, simulated, and analyzed a general model for polymeric shells. We study the response to tensile point forces acting at the poles and the shape changes that the shell undergoes as it is loaded. These simulations are tested by experiments probing the force response of the nuclear lamina and further developed to study the role of the interior chromatin in the mechanical response of cell nuclei.

## MATERIALS AND METHODS

### Model and Brownian dynamics simulations

We constructed a model of a biopolymeric shell with properties similar to those of the nuclear lamina (18–20, 48) and performed Brownian dynamics simulations using custom-written C++ code. We placed *N* monomers of diameter *a* randomly on a sphere with initial radius 2*R* and relaxed the structure for *τ*_1_ = 100 by allowing subunits to repel each other via a soft potential modeling excluded volume interactions, while remaining confined to a shell of the prescribed radius. Excluded volume interactions were modeled by the potential:

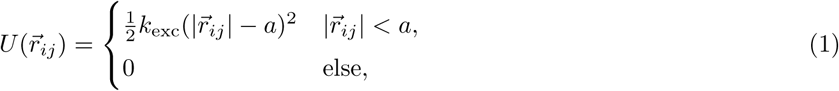

where 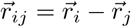 We typically take *k*_exc_ = 100*f*_0_*/a*, where *f*_0_ is the simulation force unit. During the first half of the relaxation, the prescribed radius of the shell was reduced from 2*R* to *R* at a fixed rate; during the second half of the relaxation the radius remained fixed at *R*. Except where specified in the text below, after relaxation, each subunit was connected to at least *z* = 6, but no more than *z*_max_ = 8, nearest neighbors (⟨*z*⟩ ≈ 6.5) via springs of stiffness *k*_bond_ and rest length equal to the (initial) distance between the monomers. Additional connections were made, as necessary, to ensure connectivity of the entire shell. These bonds were permanent, so the topology of the network remained fixed. To stretch the shell, individual monomers at each of the two opposite poles, identified as the two extremal monomers along the *x* axis, were pulled outward along the axis. In the simulation, this consists of applying a constant force with magnitude *F* to each of the extremal monomers, so that the shell is held at total tension *F*. For Brownian dynamics, this amounts to displacing each monomer by *F* Δ*t/ζ* at each timestep. This ensures that the force applied to each extremal monomer is *F* (in opposite directions), and that the mean tension in the polymer shell is also *F*.

To reduce computation time, we evolved the structure without Langevin noise and with lower monomer drag coefficients (henceforth referred to as “zero temperature”) for *τ*_2_ = 1000 before evolving the structure at the desired finite temperature (and corresponding monomer drag coefficients) for at least *τ*_sim_ = 7500, after which the extension of the shell was measured. We verified that this procedure did not alter our results by comparing to simulations of test cases at finite temperatures in which shells were evolved at a single fixed temperature until achieving thermal equilibrium and steady mean extension. The reported strains are averages after relaxation of at least 11 different initial configurations.

For simulations modeling a cell nucleus containing chromatin, an interior polymer was placed inside the shell at the beginning of the simulation. The shell in these simulations had *N* = 1000 subunits and initial radius *R* = 7.1*a*. Each subunit in the shell was connected to at least *z* = 4, but no more than *z*_max_ = 8, neighboring subunits by springs with stiffness *k*_bond_. The polymer was composed of *N*_p_ = 552 subunits of diameter 0.8*a*, and the subunits were connected by springs with stiffness *k*_p_ = 2*k*_bond_ to form a linear chain. At the beginning of the simulation, before relaxation, the linear chain was configured as a random walk confined within the shell. All polymer subunits repelled each other by a soft repulsive potential (modeling excluded volume interactions), which was given by Eq. 1, but with stiffness *k*_exc,p_ = *k*_p_ and rest length 0.8*a*. Soft repulsive interactions between polymeric and shell subunits were modeled by the same potential, but with stiffness *k*_exc,inter_ = *k*_exc_*k*_exc,p_*/*(*k*_exc_ + *k*_exc,p_). Of the polymer subunits, 2*N*_c_ = 110 of them were crosslinked to another polymer subunit that resided at least 4 subunits away along the polymer chain. The crosslinks were modeled by springs with stiffness *k*_c_ = *k*_p_. In addition, *N*_s_ = 40 subunits in the polymer were linked to shell subunits by springs with stiffness *k*_s_ = *k*_p_.

Simulation parameters are determined from experiments as described in (19). For simulations designed to quantitatively reproduce experiments, we take *a* = 0.7 *µ*m and *T* = 300 K. This fixes the simulation force unit to be *f*_0_ = 5.9 pN. The typical inter-subunit bond spring constant is *k*_bond_ = 100*f*_0_*/a* = 0.8 nN*/µ*m. Although we do not directly study shell dynamics, a simulation time unit corresponds to *τ ≈* 1 *µ*s.

Subunits were subject to thermal forces, 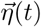, and the system was evolved by integrating the coupled Langevin equations of motion for the described interaction potentials, 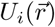, with an Euler algorithm (49) and time step Δ*t* = 0.0005. For example, in simulations of the shell alone, we integrate:

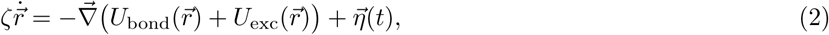

where 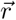 is the vector containing the 3*N* monomer position coordinates.

### Nucleus micromanipulation experiments

Individual cell nuclei were isolated from HeLa cells expressing GFP-Lamin A and mouse embryonic fibroblast cells null for vimentin (MEF V -/-) following the procedure described in (19). As discussed in (19), MEF V -/-cells are used for simpler isolation of nuclei, but the mechanical response of V -/-cells is similar to that of wild-type cells.Cells were treated with 1 *µ*g*/*mL latrunculin A (Enzo Life Sciences) for *∼* 45 minutes before single nucleus isolation.Application of 0.05% Triton X-100 in phosphate-buffered saline (PBS, Lonza) via gentle spray from an “isolation”micropipette was used to lyse the cell membrane. A second “pull” micropipette filled with PBS was used to capture the isolated nucleus by application of minimal suction pressure of *∼* 1 kPa, after which the nucleus was held by nonspecific binding between the nucleus and the interior of the micropipette.

Force-extension measurements were performed on MEF V -/-cells as in (19). The cell nucleus was held by another “force” micropipette, which was held stationary while the “pull” pipette was moved to stretch the nucleus. The speed of the pull pipette was 50 nm*/*s, but we note that nuclear mechanical response is insensitive to pulling speed over the range 15-50 nm*/*s (19). The resulting deflection of the “force” pipette, which was precalibrated as described in (50), reported the force exerted on the micropipette. To treat nuclei with MNase in force measurement experiments, another “spray” pipette was vacuum filled with 20-50 *µ*L of 1 U*/µ*L MNase solution. The solution was sprayed on the nucleus for 30-60 seconds. Measurements of nuclear spring constants were performed for 18 untreated nuclei and 5 MNase-treated nuclei.

To treat HeLa nuclei expressing GFP-lamin A with MNase in experiments in which nuclei were stretched and imaged but force response was not measured, MNase was diluted in the imaging volume at 2 U*/µ*L. Nuclei were imaged after 5 minutes and stretched beyond ≈ 30% strain. 7 nuclei were imaged and stretched in each type of experiment. For each nucleus, a 5-pixels-wide line scan (1 pixel = 90 nm) of GFP-lamin A signal was measured through the center of the nucleus, perpendicular to the stretching axis. We used a custom-written Python code to identify peaks in GFP-lamin A signal. Briefly, the data was smoothed and outermost local maxima in signal intensity were identified as the peaks marking the boundaries of the nucleus. Between boundary peaks, fluorescence peaks exceeding the mean signal by at least 15% were counted as peaks that indicate the presence of axial buckles.

## RESULTS

### Biopolymeric shell model

We constructed a simulation model of a biopolymeric shell with properties similar to those of the nuclear lamina (18–20, 48). We placed *N* monomers of diameter *a* randomly on a sphere of radius *R* and relaxed the structure by allowing subunits to repel each other via a soft repulsive potential (modeling excluded volume interactions) with stiffness *k*_exc_ (see Eq. 1), while remaining confined to a shell of the prescribed radius. Each subunit was connected to at least *z* = 6 nearest neighbors (⟨*z*⟩ ≈ 6.5) via springs of stiffness *k*_bond_ and rest length equal to the distance between the monomers in the initial shell configuration. The structure is a single randomly linked polymeric layer with fixed network topology, comprising a spherical shell with weak bending elasticity that models the structure and microscopic mechanics of the lamin A/C intermediate filament network of the nuclear lamina (17, 18, 20, 48, 51, 52). To probe the mechanical response of the model in an experimentally testable manner, we conducted Brownian dynamics simulations of the structure, and we held the shell at total tension *F*.

### Biopolymeric shell model exhibits a two-regime mechanical response

These biopolymeric shells generically exhibit a two-regime mechanical response (Fig. 1A). At zero tension, shells have mean diameter ⟨*L*⟩ ≡ *L* ≈ 2*R*. They exhibit linear response to small tensions but stiffen beyond extensions, Δ*L*, of order *R*, and cross over into a large-tension regime of a stiffer, but still linear, response (Fig. 1A). Since the shell is composed of purely linear springs, this crossover is due to the geometry of the shell. As the shell deforms, an increasing number of the springs in the shell align with the tension axis, and thus, the shell resists extension more strongly.

**Figure 1:**
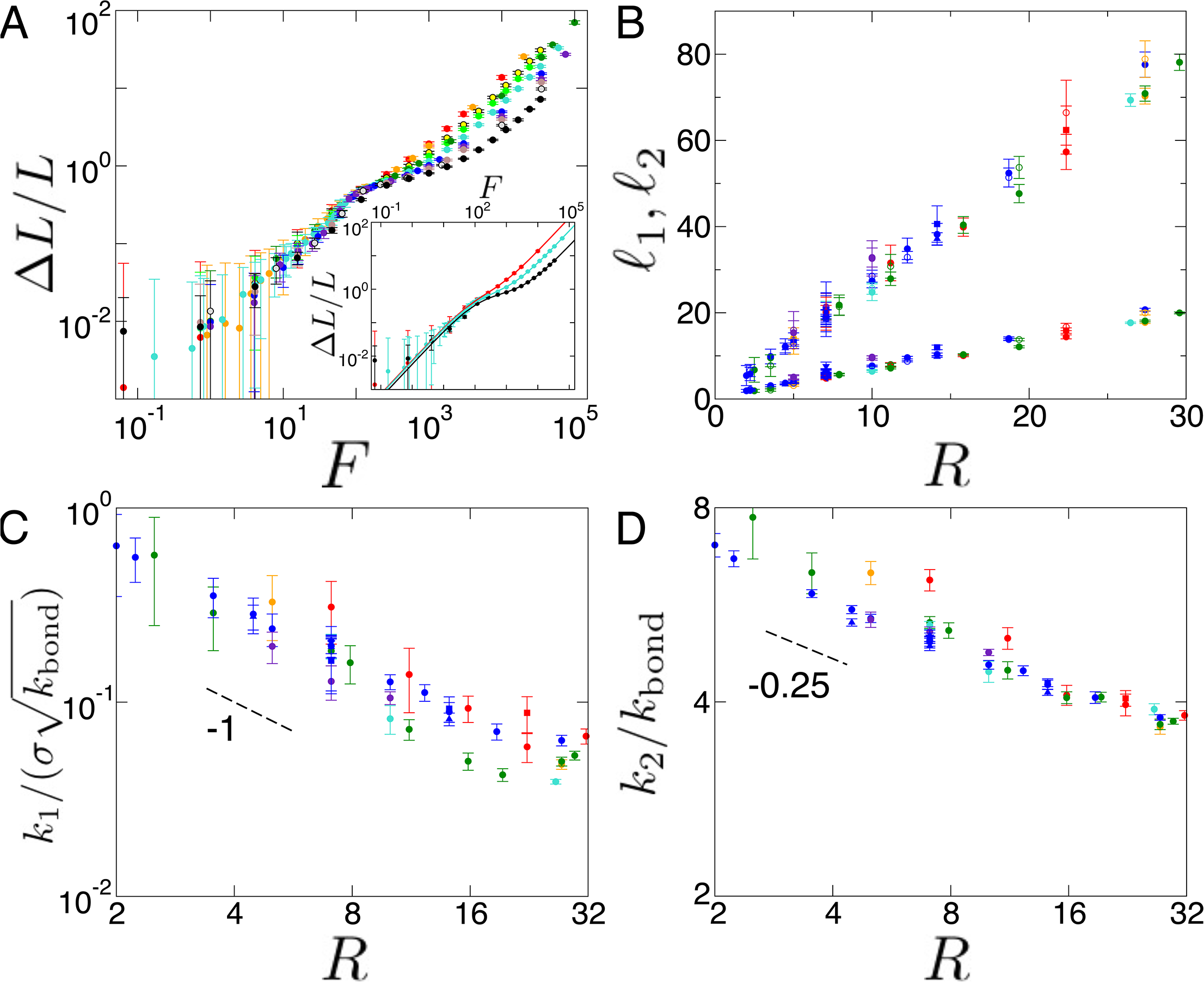
Tension-strain relation for polymeric shells. (A) Shell stretched by tension *F* exhibits two-regime response, with linear response at small tensions and stiffer linear response to large tensions. Tension-strain curves do not scale simply with *N* (σ ≡ *N/R*^2^ = 20 with *N* = 80, red; 100, orange; 250, yellow; 400, light green; 500, dark green; 1000, turquoise; 2000, blue; 3000, purple, 4000, brown; 7000, gray; 15000, black). Inset: Examples of fitting by Eq. 3. (B) Crossover lengths, 𝓁_1_ and 𝓁_2_, scale linearly with shell radius, *R*. Colors, shapes, and filling of symbols indicate different *σ*, *k*, and *k*_*B*_*T*, respectively (σ = 2, red; 4, orange; 8, dark green; 10, turquoise; 20, blue; 40, purple. *k* = 10, downward pointing triangle; 25, square; 50, diamond; 100, circle; 200, triangle up; 400, triangle left. *k*_*B*_*T* = 10^−3^, filled; 1, open). (C) Small extension spring constant scales as 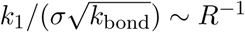. (D) Large extension spring constant scales as *k*_2_*/k*_bond_ *∼ R*^−0.25^. Symbols for C and D are as in B. Lengths are in units of *a*, tensions in simulation force units of *f*_0_, spring constants are in units of *f*_0_*/a*, and temperatures in units of *f*_0_*a*. Measurements are from at least 11 simulations per data point.

Empirically, the tension-strain relation can be fit by (inset to Fig. 1A):

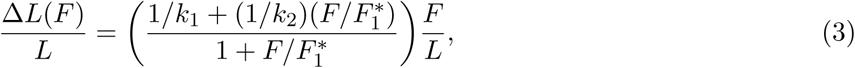

which has linear tension-extension relations in the limits of small and large tensions, with respective spring constants *k*_1_ and *k*_2_. Eq. 3 defines two crossover force scales, 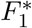 and 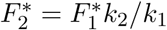, and correspondingly, two crossover length scales:

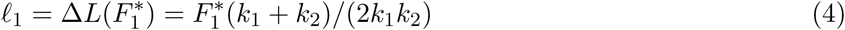

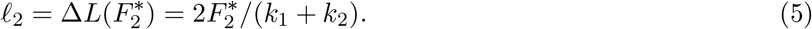

The crossover lengths scale linearly with the radius, *R*, of the shell, and they are insensitive to number of subunits in the shell *N*, bond stiffness *k*_bond_, and temperature *k*_*B*_*T* (Fig. 1B). Since *k*_2_ ≫ *k*_1_, 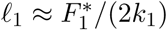and 𝓁_2_*/*𝓁_1_ *≈* 4.

The simulation tension-strain relation does not obey a single simple scaling law (Fig. 1A), but the spring constants *k*_1_ and *k*_2_ scale with *R*. The small-tension spring constant scales as

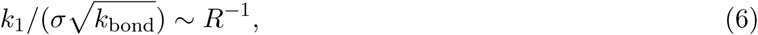

where *σ ≡ N/R*^2^ is the surface density of monomeric subunits in the shell (Fig. 1C). *k*_1_ *∼ σ* arises from the local stiffening to small deformations that accompanies denser polymer networks. The *R*^−1^ scaling factor is expected based on linear elasticity theory (38) and is also reported for experiments with polymerized vesicles (12). The deformation over a region of area *d*^2^ due to force *F* costs bending energy *κ*Δ*L*^2^*/d*^2^ and stretching energy *k*_*s*_(Δ*L/R*)^2^*d*^2^, where *κ* is the emergent bending modulus of the shell and *k*_*s*_ is the spring constant for local stretching. By minimizing the energy with respect to *d*, we find *d ∼* (*κR*^2^*/k*_*s*_)^1*/*4^. Setting these two energy terms to equal the work, *F* Δ*L*, done by the force, we find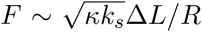. Since the inter-subunit bond stiffness controls the shell stretching stiffness, we have 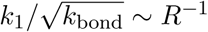

For large tensions, the shell spring constant scales as (Fig. 1D)

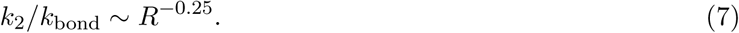

This deviates from the scaling of *k*_2_ *∼ R*^0^ that one might expect for a two-dimensional spring network. The expectation is that for a 1D lattice of length *R* and line density *λ* of springs, the total spring constant is *K ∼* (*λR*)^−1^; for a 2D *R × R* lattice with spring density *σ*,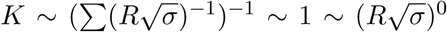; in 3D, *K ∼ ρ*^1*/*3^*R*. Qualitatively, the observed scaling for *k*_2_*/k*_bond_ arises because the tension deforms the spherical shell primarily in one dimension, which reorients the springs along the tension axis and effectively lowers the dimensionality of the network. This scaling appears paradoxical since it suggests that *k*_2_ *→* 0 for a flat polymerized membrane (*R → ∞*). However, in that case, 𝓁_2_ *→ ∞*, so the large-tension linear regime does not exist, and there is no inconsistency. *k*_2_ *∼ k*_bond_ because shell deformations directly stretch inter-subunit springs in the large-tension regime.

The simulations thus reveal two distinct scaling regimes of polymeric shell elasticity arising from the geometry of the spherical shell and axial deformation. The corresponding spring constants, *k*_1_ and *k*_2_, are defined by the network density, bond stiffness, and shell size, while the crossover length scales, 𝓁_1_ and 𝓁_2_, depend sensitively on only the size of the shell.

### Shells under tension undergo an azimuthal-symmetry-breaking buckling transition

When the polymeric shell is extended by *≈* 𝓁_1_, it undergoes a buckling transition (Fig. 2A), similar to those of flat elastic membranes (53–55), elastic shells (44, 47, 56), and flat polymerized membranes (43). As the shell is extended along the tension axis, it contracts along its transverse axis (Fig. 2B), which exerts circumferential compressive stresses that cannot be relieved due to the topology of the shell. For sufficiently high compression, it is energetically favorable to relieve these compressive stresses through localized bending (43, 53–55), which manifests in the polymeric shell as some subunits shifting out-of-plane with relation to neighboring subunits. The cost in deformation energy is minimized by aligning nearby bends, which leads to the formation of buckles that extend from pole to pole.

**Figure 2:**
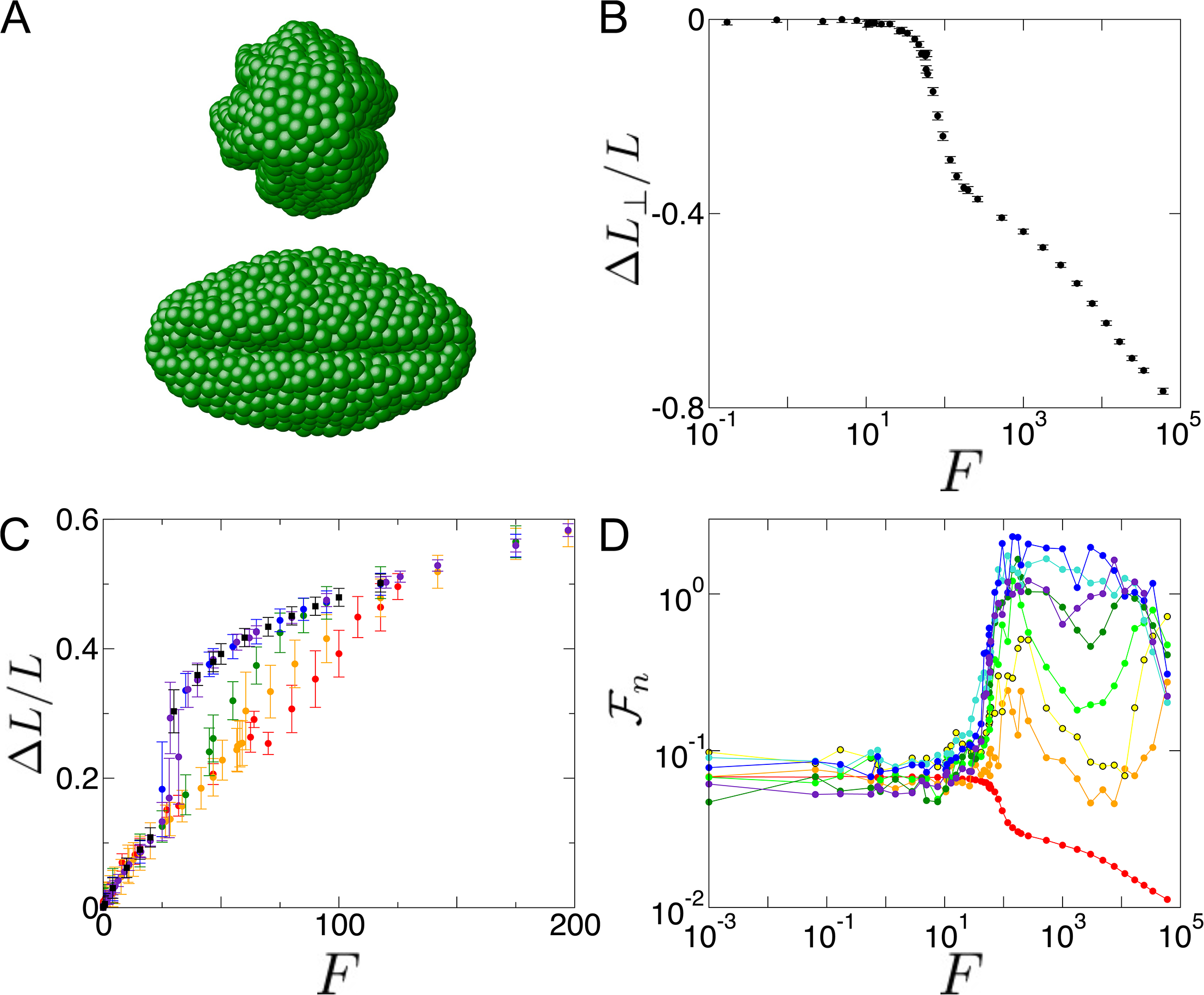
Buckling of stretched polymeric shells. (A) Two views of stretched shell with multiple buckles perpendicular to tension axis extending longitudinally across the shell. (B) Transverse strain, Δ*L*_*⊥*_*/L*, versus tension, *F*, drops sharply at the buckling transition. (C) Tension-strain relation exhibits signature of buckling transition, with a jump in strain that increases with increasing temperature (*k*_*B*_*T* = 0 [no Langevin noise, see Materials and Methods], red; 10^−3^, orange; 10^−2^, green; 10^−1^, blue; 1, purple). Black squares show hysteretic behavior of force-strain relation for shells evolved at *k*_*B*_*T* = 10^−3^ after a transient of *k*_*B*_*T* = 1. (D) Normalized Fourier modes, 𝔉_*n*_(*F*) (Eqs. 8-9), indicate buckling by increases in modes with *n ≥* 2 at tensions corresponding to the jump in the tension-strain curve. *n* = 0, red; 2, orange; 3, yellow; 4, light green; 5, dark green; 6, turquoise; 7, blue; 8, purple.

Buckling manifests itself in the tension-strain curves as a temperature-dependent jump at moderate tensions (Fig. 2C). Equivalently, this can be viewed as a stiffness-dependent jump; since the spring constant in the simulation is *k*_bond_ = *κ*_0_*f*_0_*/a* and the temperature is *k*_*B*_*T* = *τ*_0_*f*_0_*a*, temperature and stiffness are related by 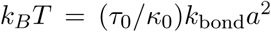, where *κ*_0_ and *τ*_0_ are dimensionless numbers. For low temperatures, the tension-strain curve is relatively smooth (red and orange in Fig. 2C), but for higher temperatures (blue and purple), the jump in the tension-strain curve is *≈* 20% in strain. Alternatively, considering force as a function of strain, the jump can be interpreted as a plateau in the tension, which is suggestive of phase coexistence. Moreover, the shell loading curve exhibits hysteresis; stretched, buckled shells from high temperature simulations remain in the larger extension state after *T* is decreased (black in Fig. 2C). Altogether, this suggests a first-order phase transition, paralleling the behavior of pressurized shells (46, 47).

We quantify buckling by defining and measuring the mean normalized Fourier modes as a function of tension:

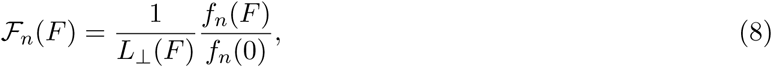

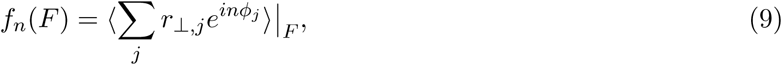

where *L*_*⊥*_(*F*) is the mean transverse shell diameter as a function of tension, *r*_*⊥,j*_ is the distance of the *j*^th^ subunit from the line parallel to the tension axis running through the shell center of mass, *φ*_*j*_ is the azimuthal angular coordinate of the *j*^th^ monomer, and the summation is performed over all monomers in the shell. The zero-tension amplitudes scale as *f*_*n*_(0) *∼ n*^−1^ in the simulation. At the jump in the tension-strain curve, *L*_*⊥*_(*F*) and 𝔉_0_(*F*) drop, Fourier modes with *n ≥* 2 increase sharply (Fig. 2D), and the shell buckles into a multi-lobed structure. This is again indicative of a first-order phase transition, which is expected from the symmetry of these modes. This can also be seen by constructing a Landau theory with rotationally invariant combinations of the *F*_*n*_, similar to what is done with spherical harmonics in (47) for shells under pressure.

### Buckling can be observed for dechromatinized nuclear laminas

We investigated whether the morphology predicted by the simulation can be experimentally observed for a biological structure. We considered the nuclear lamina, which is a polymeric mesh shell of intermediate filaments with mesh size *∼* 0.4 *µ*m (17–20). To study the lamina, we isolated individual HeLa and MEF V -/- cell nuclei following the procedure described in (19) (see Materials and Methods). Upon isolation, nuclei were treated with the enzyme MNase, which fragments the chromatin contained within the nucleus; fragmented chromatin exits the nucleus, leaving behind an isolated lamina. Based on previous quantitation by Hoechst staining, *<* 40% of chromatin content remains in the nucleus after MNase treatment (19). Forces were applied by two micropipettes attached to opposite poles of the lamina and steadily separated via micromanipulation apparatus. Deflection of the micropipettes upon nuclear stretching reports the force, which is typically of order nN (19). Thus, the geometry and force application is similar to that of our model, but distinct from previous micropipette aspiration measurements performed on nuclei (16, 17, 22–24), red blood cells (15, 57, 58), and polymer and fluid vesicles (59, 60).

The nuclear lamina exhibits mechanics and morphology that are consistent with the polymeric shell model. At zero applied force, the lamina fluctuates but maintains its round, inflated shape, as expected for polymeric shells (45, 61, 62) (Fig. 3A, left). In response to small applied force, the lamina extends with a soft spring constant of order 0.1 nN*/µ*m. However, as extension approaches the nuclear radius, the lamina stiffens to *∼* 1 nN*/µ*m (Fig. 4A).

**Figure 3:**
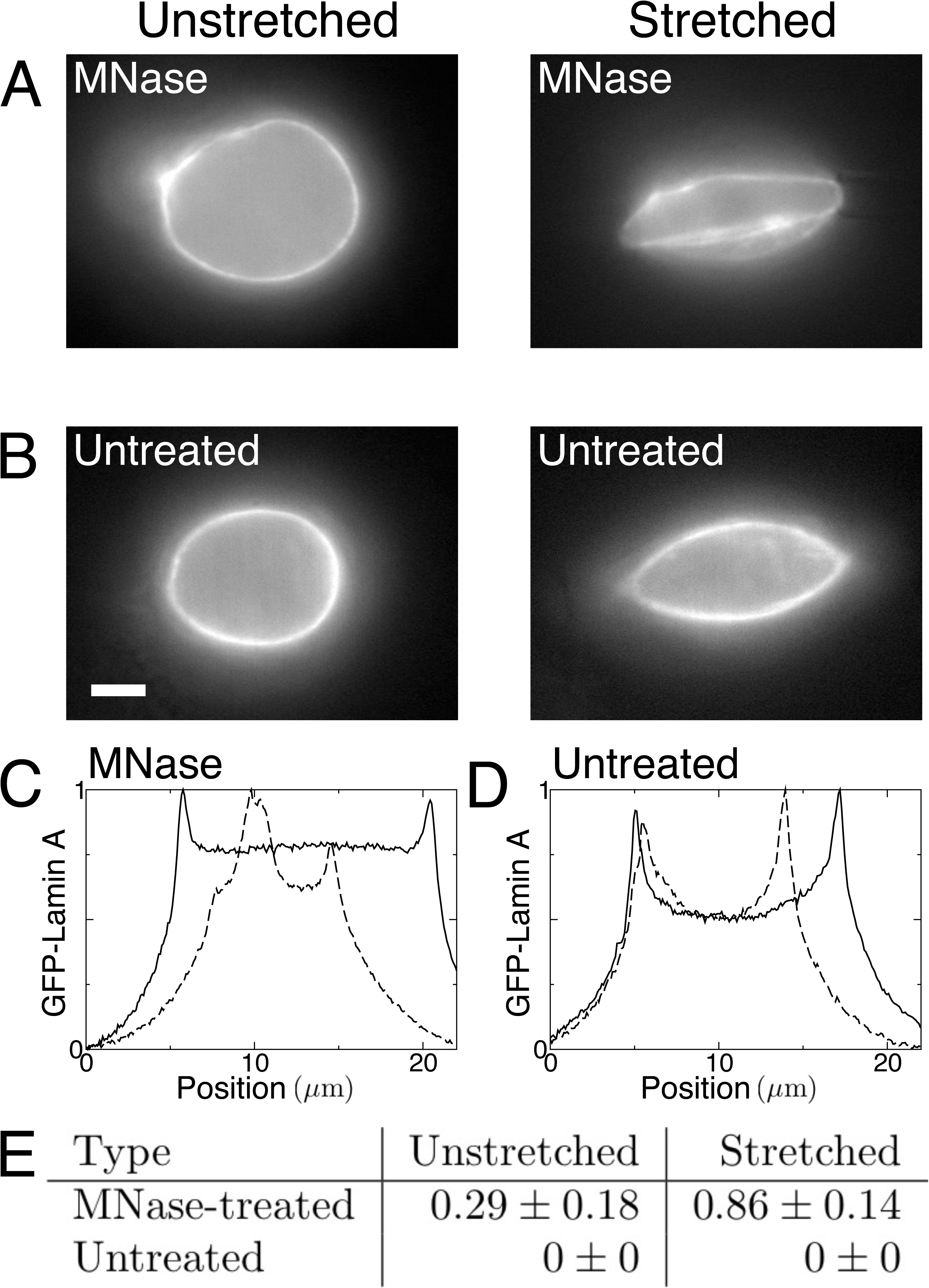
Stretched laminas, but not the nuclei containing chromatin, exhibit axial buckles. (A) Representative images of an unstretched (left) and stretched (right) lamina, obtained by treating HeLa nuclei with MNase, which digests chromatin. Stretched image shows longitudinal buckles. (B) Representative images of nuclei with intact chromatin interior, which do not exhibit buckling when stretched (right; compare to left, unstretched). Scale bar is 5 *µ*m. Fluorescence signal is GFP-lamin A. (C) Line scans along the central axis perpendicular to the tension axis showing the GFP-lamin A signal for an MNase-treated nucleus when unstretched (solid line) and stretched (dashed line), corresponding to the images in A. GFP-lamin A intensity is plotted in arbitrary units. (D) Line scans along the central axis perpendicular to the tension axis showing the GFP-lamin A signal for an untreated nucleus when unstretched (solid line) and a stretched (dashed line), corresponding to the images in B. (E) Table listing average number of excess peaks (between boundary peaks) per line scan. 7 nuclei were imaged for each case. Error is given by standard error of the mean.

**Figure 4:**
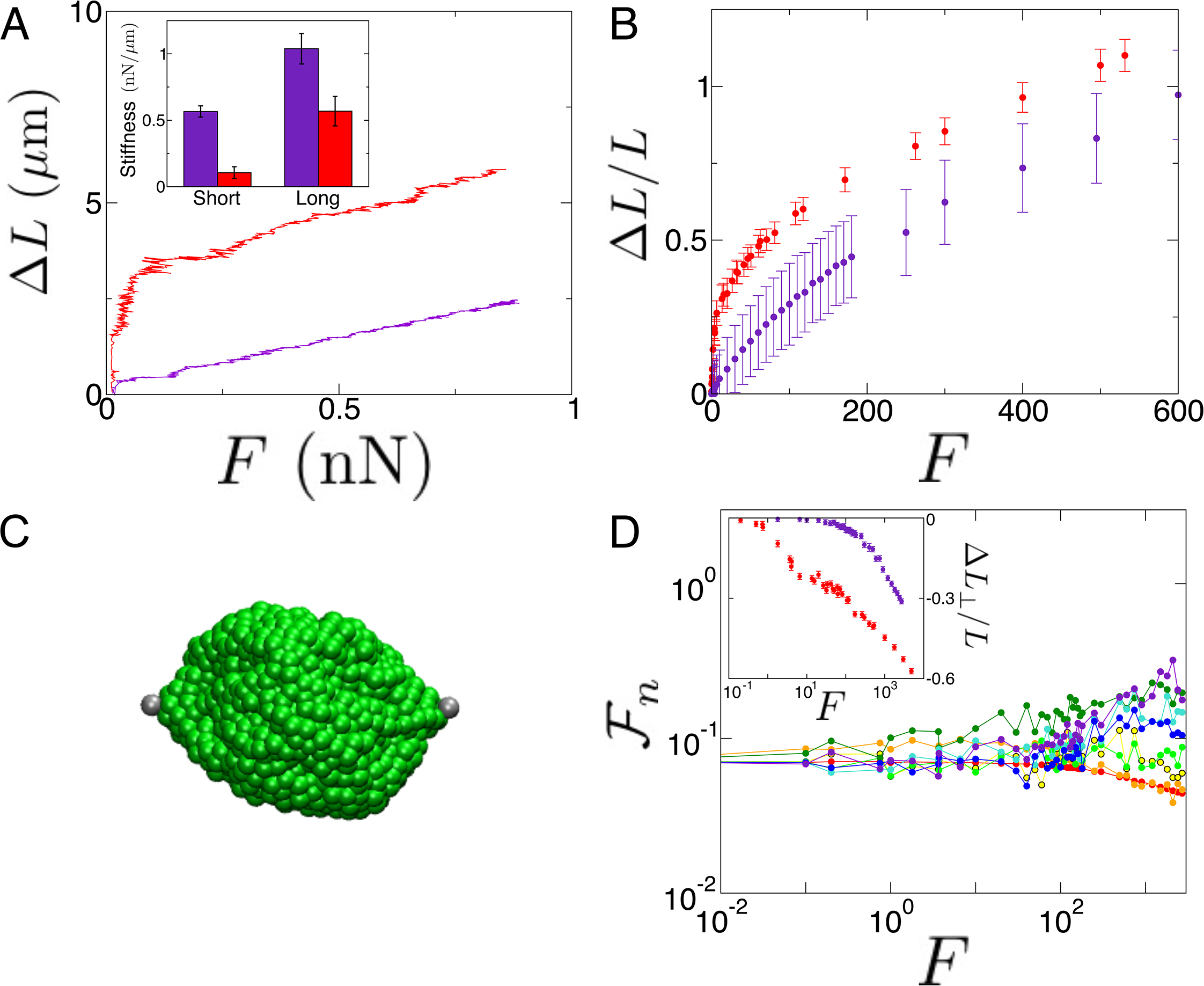
Shells filled with a tethered crosslinked polymer, modeling a cell nucleus, do not axially buckle. (A) Force-extension curves for MEF V -/- nuclei in experiments in which nuclei were treated with MNase (red) and without (purple) MNase treatment. Inset: Bar graph showing effective spring constants for MEF V -/- nuclei in short- and long-extension regimes with (red) and without (purple) MNase treatment. Measurements were performed for 18 untreated and 5 MNase-treated nuclei. Error bars are standard error of the mean. Statistical significance of *p <* 0.05 was established by T-test. (B) Tension-strain relation shows that the tethered crosslinked polymer interior strengthens initial force response (purple, as compared to empty shell, red). Simulation units are *a* = 0.7 *µ*m and *f*_0_ = 5.9 pN. (C) While there are localized buckles in the stretched shell with a tethered polymer interior, buckles do not span the entire structure. (D) Normalized Fourier modes show weaker signature of buckling, compared to those of Fig. 2D. *n* = 0, red; 2, orange; 3, yellow; 4, light green; 5, dark green; 6, turquoise; 7, blue; 8, purple. Inset: Transverse strain for shells enclosing a tethered polymer (purple) and empty shells (red). *z* = 4 (*⟨z*⟩ *≈* 4.5) in B, C, and D.

For such deformations (≳30% strain), the lamina exhibits axial buckles that extend across the structure (Fig. 3A, right), as in the model. These buckles are observable as lines of GFP-lamin A fluorescence extending from pole to pole across the lamina. We quantify the signal of buckling in the images by performing line scans along the central axis of the lamina, perpendicular to the tension axis. As shown by representative line scans in Fig. 3C, both relaxed and extended laminas have peaks delineating the boundaries (Fig. 3C, solid and dashed lines, respectively). Between these boundary peaks, stretched nuclei/laminas typically also exhibit an additional peak whereas unstretched laminas do not. This additional peak is due to the GFP-lamin A line extending across the lamina. We quantify the set of line scans by measuring the mean number of peaks per scan. We measure an average of ⟨*n*_peaks_⟩ = 0.29 *±* 0.18 in unstretched laminas and ⟨*n*_peaks_⟩ = 0.86 ± 0.14 for stretched laminas, which demonstrates that stretched laminas consistently axially buckle, whereas relaxed laminas do not (Fig. 3E). These results are consistent with visual observations, from which it is evident that 1 of 7 relaxed laminas and 6 of 7 stretched laminas have a strong signal of axial buckling. Altogether, these results are consistent with the model prediction that biopolymeric shells undergo an axial buckling transition when deformed past a threshold extension approximately equal to the shell radius.

### Buckling is suppressed by chromatin in nuclei and a crosslinked polymer interior in the model

To further understand cell nuclear and shell mechanics and their respective buckling transitions, we performed experiments on untreated isolated cell nuclei that retained their chromatin interiors and performed simulations in which the shell was filled with a crosslinked polymer matrix. We observed that buckling and mechanics depend sensitively on the interior structure.

When chromatin is present inside the nucleus, it suppresses buckling. This is the case in nuclei not treated with MNase, where we only observe localized folds and ruffles that do not extend across the entire nucleus (Fig. 3B). This is also observed in individual GFP-lamin A line scans, which generically fluctuate about a low-signal mean without exhibiting peaks between the boundaries (Fig. 3D). Similarly, our quantitative analysis of the line scans for untreated (chromatin-filled) nuclei did not detect any peaks between the boundaries in either relaxed or stretched nuclei (Fig. 3E). These results are consistent with visual observations that 0 of 7 untreated nuclei (unstretched and stretched) exhibit axial buckling. Inhibition of buckling due to the stiff interior is also evident in nucleus micropipette aspiration experiments (22) and simulations of an elastic core-shell model (56).

In addition to its effects on morphology, chromatin also alters mechanical response of the nucleus. Untreated nuclei exhibit a stiff mechanical response to small deformations, in contrast to empty nuclear laminas (Fig. 4A). Similar to empty nuclear laminas, however, chromatin-filled nuclei stiffen in response to large deformations, primarily due to the strain-stiffening response of the nuclear lamina (19).

To model mechanical response and buckling of cell nuclei, we extended the model to capture contributions from the chromatin interior in addition to the polymeric nuclear lamina. We thus included a crosslinked polymer interior and linked it to the shell by springs. To convert from simulation to experimental units, we take *a* = 0.7 *µ*m and fix *T* = 300 K so that *f*_0_ = 5.9 pN (see Materials and Methods). For parameters based on the physical properties of cell nuclei (19), the composite shell/polymer model quantitatively captures key features of nuclear mechanical response. The presence of the tethered crosslinked polymer interior markedly stiffens the small-tension response (Fig. 4B). However, when the simulated nucleus is extended by *>* 𝓁_1_, the stiffness is dominated by the properties of the exterior polymer shell, as shown by the similar slopes for large tensions in the tension-strain relations for empty and filled shells (Fig. 4B). These results are consistent with experiments demonstrating the cell nuclear mechanical response is primarily determined by chromatin at small extensions and by the lamina at large extensions; additionally, model parameters can be modulated to mimic biophysical perturbations of cell nuclei (19).

As in the experiments with nuclei retaining chromatin, coordinated axial buckles do not appear in simulations of the composite shell/polymer model (Fig. 4C). This is confirmed by measuring the normalized Fourier modes, 𝔉_*n*_ (Eqs. 8-9), of the shell (Fig. 4D). Although there is some signature of localized buckling at moderate tensions, it is greatly suppressed relative to empty shells (Fig. 2D). Moreover, the tension-strain relation for a shell encapsulating a tethered crosslinked polymer does not display the jump that occurs for empty shells (Fig. 4B compared to Fig. 2C).

Together, the experiments and simulations demonstrate that the mechanical properties of the interior of a biopolymeric shell (chromatin in the case of cell nuclei) tightly controls both mechanical response and morphological features of the entire structure.

## DISCUSSION

We have studied a class of polymeric shells which provides a quantitative model for cell nuclear mechanical response. The model displays a novel two-regime elastic response similar to that observed in micromanipulation experiments on cell nuclei (19). At extensions of Δ*L ∼ R* (strains of *∼* 30−50%), both model shells and isolated nuclear laminas undergo a novel symmetry-breaking buckling transition with hallmarks of a phase coexistence, as occurs during a first-order phase transitions.

### Applications to nuclear mechanical response and morphology

The model quantitatively explains why cell nuclei exhibit two regimes of mechanical response dominated by two distinct structural components. For small applied forces, the lamina (shell) spring constant is small *(∼*0.1 nN*/µ*m) and depends only weakly on the microscopic (inter-node) spring constant of the lamin network 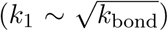. Therefore, the stiffer chromatin (crosslinked polymer) interior, which is a three-dimensional material with a Young’s modulus of order kPa (19, 24, 63, 64), determines nuclear mechanical response to small deformations. However, the nuclear lamina stiffens several-fold for extensions of order 𝓁_1_ *∼ R*, when the filaments (springs) in network align with the applied force. Moreover, the limiting spring constant, *k*_2_, scales linearly with the microscopic spring constant, *k*_bond_. Thus, the lamina’s (shell’s) mechanical properties are dominant in nuclear response to large applied forces, and the transition between the two regimes of mechanical response may be governed by the spheroidal geometry of the nucleus.

In addition to these mechanics, the appearance of length-spanning buckles in the empty shell model and the isolated nuclear lamina suggests an important role for the chromatin interior in maintaining nuclear morphology. Because the nucleus is continually subjected to intra-and extracellular forces *in vivo*, it must be able to maintain its shape against perturbations. Since buckling is hysteretic for shells with the mechanical properties of cell nuclei (19, 47) (Fig. 2C), large deformations would not relax and could thus permanently disrupt the nucleus. However, chromatin remedies this problem by providing a stiff scaffold that prevents formation of length-spanning buckles (Figs. 3B and 4). This is consistent with previous work showing that buckling morphology of spheroidal shells can be controlled by the stiffness of the interior (56, 65). Thus, an interesting open question is to what extent chromatin stiffness dictates nuclear morphology, as compared to the structural and compositional perturbations of the nuclear lamina which have been explored in previous models (33, 34).

### Physics of the buckling transition

The first-order nature of the buckling transition can be understood by considering the form of the free energy for the shell in terms of azimuthal order parameters (the Fourier amplitudes plotted in Fig. 2D). If we associate an order parameter *a*_*n*_ with each Fourier mode *f*_*n*_ (Eq. 9), and note that it corresponds to an azimuthal density *∝ e*^*inφ*^ where *φ* is the azimuthal angle, we immediately see that there will be quadratic terms in the free energy proportional to *|a*_*n*_*|*2 for all *n*, but also that there will be *cubic* terms. The lowest order nontrivial cubic terms in the free energy are 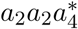and 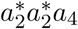(*a*_1_ is absent since it corresponds to a simple global translation in the *x*-*y* plane). These cubic terms in the free energy guarantee that the transition will be first-order in character in much the same way as occurs in the density-wave theory of crystallization (66) or in isotropic-to-nematic liquid crystal phase transitions (67). Similar arguments have also been made for symmetry breaking during compressive buckling of pressurized elastic shells (47).

### Comparison with previous models for cell nuclei

Several models have been developed previously to describe nuclear mechanics and morphology, but several major features of our biopolymeric shell model contrast with these models (17, 23, 33, 34, 68). First, the biopolymeric shell is composed of discrete subunits, in contrast to previous continuum models (17, 23, 33, 68). This feature facilitated the study of the dependence of shell mechanics on number, *N*, of subunits and allowed the inter-subunit connectivity, *z*, to be altered as desired (Fig. 1). Moreover, both of these variables have a clear corresponding meaning in real nuclear laminas. Secondly, the subunits softly repel each other, in contrast to a previous spring network model (34). This additional assumption leads to the clear axial buckling that we observe when the shell is stretched (Fig. 2). Third, in contrast to the continuum models (17, 23, 33, 68), our model does not include intrinsic bending stiffness or spontaneous curvature. This assumption also enhances buckling, and it is consistent with the properties of lamins and other intermediate filaments (20, 48, 52). Fourth, inter-subunit bonds are permanent and subunits do not diffuse within the network, and fifth, subunits are arranged in a single layer. In these two aspects, two previous models (33, 34) may more closely realize the experimental situation for cell nuclei, in which lamin filaments can rupture and lamin proteins can exchange (21, 69) and lamins A and B form distinct, but connected networks (69). We predict that including inter-subunit bond dynamics would modulate the viscoelastic behavior of the shell and possibly decrease the observed effective spring constants. Nonetheless, our more minimalistic approach has the benefit of modeling the behavior of a broader class of biopolymeric shells.

### Nuclear mechanical response *in vivo*

While our model describes the experimentally observed mechanical response of nuclei, it does not include possible contributions from the cellular environment in which the nucleus resides. The cell is a mechanically active environment, and its cytoskeleton continually deforms and reorganizes via actin polymerization and depolymerization and myosin-driven contractions. The resulting stresses could be transmitted to the nucleus via actin stress fibers comprising the perinuclear actin cap, which are connected to the nucleus via LINC complexes in the nuclear envelope (70). In general, such “prestresses” can stiffen the mechanical response and alter the dynamics of semiflexible polymer and actin networks (71–76). Our model typically includes some degree of prestress due to overlaps between monomeric subunits when the shell is constructed. However, in principle, additional prestress or active stresses could imbue the nuclear lamina with additional stiffness in response to small deformations by tensing the lamin network.

Nonetheless, the effects of these active prestresses and other environmental factors on *in vivo* nuclear mechanical response is unclear. Previous micropipette aspiration (17) and micromanipulation experiments (19) performed on nuclei remaining in cells found little difference between the mechanical response of *in vivo* nuclei and that of isolated nuclei. Additional micromanipulation experiments found that nuclear deformability is not affected by depolymerization of the actin cytoskeleton, but is increased by loss of the intermediate filament vimentin, which may provide mechanical support by caging the nucleus while it remains in the cell (26). Such effects could be incorporated into a refined version of our biopolymeric shell model.

### Conclusion

Altogether, the simulations provide an experimentally testable, biologically relevant, and quantitatively predictive model for the mechanics of biopolymeric shells. These shells have two linear tension-strain regimes, corresponding to a weak, linear elastic response to small tensions and a stiff linear response when the shell deforms sufficiently to align the inter-subunit bonds with the tension axis. The small-tension spring constant scales as *k*_1_ *∼ R*^−1^, as expected from linear elasticity, but surprisingly, the large-tension spring constant scales as *k*_2_ *∼ R*^−0.25^, insensitive to *N*. Between these regimes, the shell undergoes a buckling transition in which multiple length-spanning folds appear. Buckling can be inhibited by a stiff interior, such as a crosslinked polymer linked the shell. Both strain stiffening and the axial buckling can be experimentally observed via micromanipulation of isolated cell nuclei. These observations highlight the importance of the chromatin interior in cell nuclear mechanical response and shape maintenance. More generally, the model could provide insights into the behavior of a wider class of biopolymeric shells such as protein-coated vesicles (*e.g.*, actin or clathrin (5–12)) and viral capsids (2, 3).

## AUTHOR CONTRIBUTIONS

EJB, ADS, and JFM designed research and discussed results, EJB performed simulations and theoretical calculations, ADS performed experiments, EJB and ADS analyzed data, and EJB wrote the manuscript.

## ACKNOWLEDGMENTS

We thank R. D. Goldman and S. A. Adam for helpful discussions and A. Erbaş and S. Brahmachari for discussions and critically reading the manuscript. We acknowledge support by NSF Grants MCB-1022117 and DMR-1206868, by NIH Grants 1U54CA193419 and 1R01GM105847, and by subcontract to the University of Massachusetts under NIH grant U54DK107980. A.D.S. also acknowledges support by NRSA postdoctoral fellowship F32GM112422 and by a postdoctoral fellowship from the American Heart Association 14POST20490209 (7/1/14 - 2/29/16). This research was supported in part through the computational resources and staff contributions provided for the Quest high performance computing facility at Northwestern University, which is jointly supported by the Office of the Provost, the Office for Research, and Northwestern University Information Technology.

